# Centrioles are amplified via rosette formation in cycling progenitors of olfactory sensory neurons

**DOI:** 10.1101/695098

**Authors:** Kaitlin Ching, Tim Stearns

**Affiliations:** Department of Biology, Stanford University, Stanford, CA 94305, USA; Department of Genetics, Stanford University School of Medicine, Stanford, CA 94305, USA

**Keywords:** olfaction, centriole, cilium, OSN, stem cell, single-cell RNAseq

## Abstract

Olfaction in most animals is mediated by neurons bearing cilia that are accessible to the environment. Olfactory sensory neurons (OSNs) in chordates usually have multiple cilia, each with a centriole at its base. OSNs differentiate from stem cells in the olfactory epithelium, and how the epithelium generates cells with many centrioles—about 16 in mouse—is not yet understood. We show that centrioles are amplified via centriole rosette formation in both embryonic development and turnover of the olfactory epithelium in adults, and rosette-bearing cells often have free centrioles in addition. Cells with amplified centrioles can go on to divide, with centrioles clustered at each pole. Additionally, we found that immediate neuronal precursors amplify centrioles concomitantly with elevation of mRNA for Plk4 and Stil, key regulators of centriole duplication. Our findings highlight the importance of accounting for centriole amplification in neuron regeneration therapies derived from olfactory epithelia.

## Introduction

Olfaction, the primary way that animals sense their chemical environment, begins in olfactory sensory neurons (OSNs). In many chordates, each OSN has multiple cilia which protrude from the end of a dendrite at the apical surface of the olfactory epithelium. At the apical surface, odorants contact receptors on the surface of cilia, initiating a signaling event in the OSN. At the base of each cilium, a centriole organizes the structure (**Figure 1A**). Cilia are necessary for olfaction, as are the centrioles which organize their microtubule structures. To have multiple cilia, each OSN must have multiple centrioles, raising the question: how are these many centrioles made?

**Fig. 1.**
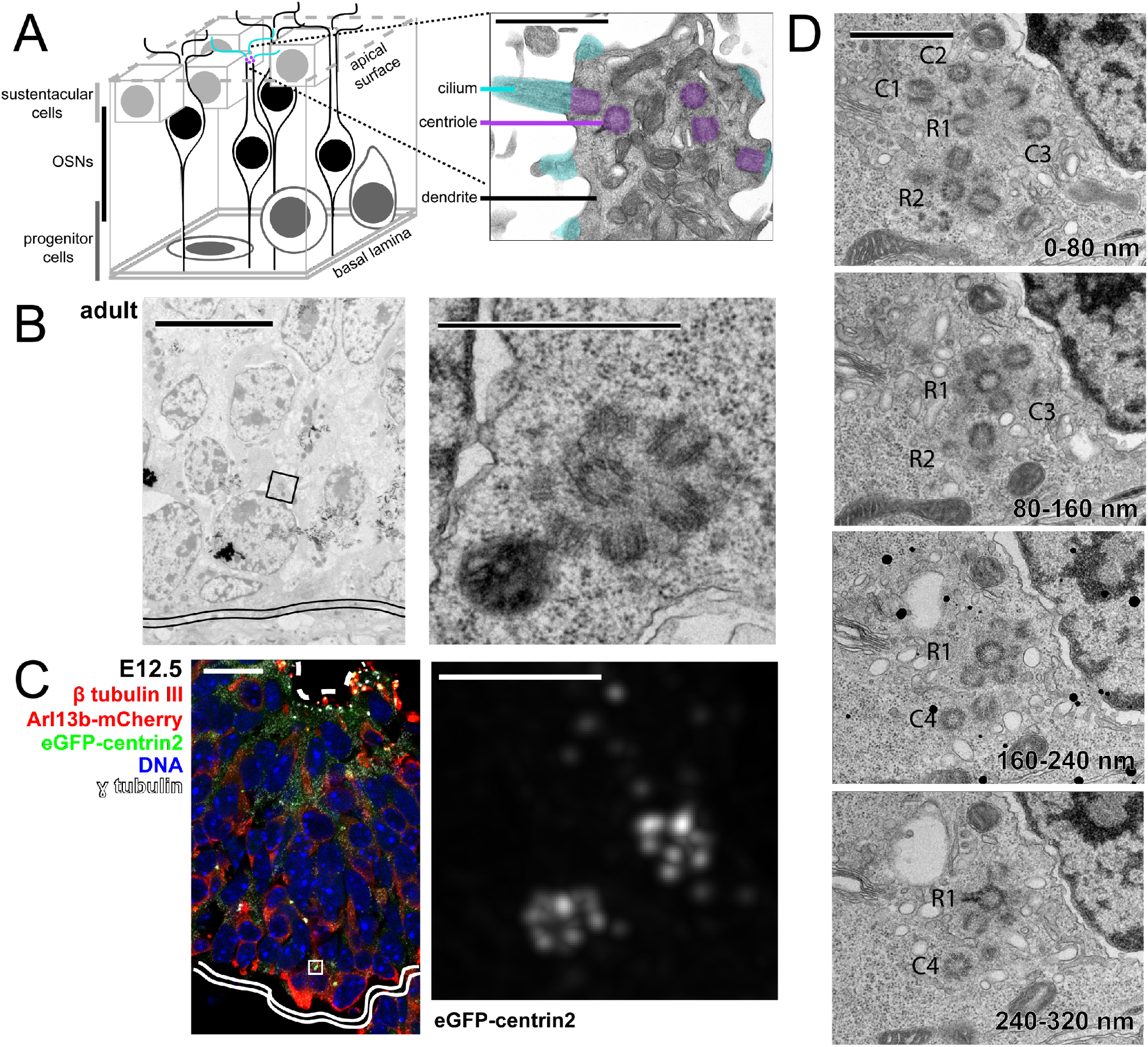
Centriole locations and rosette structures in adult and embryonic mouse olfactory epithelium. (A) Schematic of the olfactory epithelium. The inset shows an OSN dendrite from adult mouse imaged by TEM with pseudocolored cilia and centrioles. Inset scale bar = 1 μm. (B) TEM image of wild-type adult mouse olfactory epithelium. The inset shows a centriole rosette in cross section, near the basal lamina. Double solid line marks the basal lamina. Scale bar = 10 μm. Inset scale bar = 1 μm. (C) Fluorescence image of embryonic olfactory epithelium at E12.5 in mice expressing eGFP-centrin2 to mark centrioles. The inset shows a deconvolved image of two rosette-like centriole clusters from a cell positive for β tubulin III, near the basal lamina. Dashed line marks the apical surface of the olfactory epithelium. Double solid line marks the basal lamina. Scale bar = 20 μm. Inset scale bar = 2 μm. Also see figure S1A. (D) TEM images of wild-type adult mouse olfactory epithelium in serial sections. The images show two centriole rosettes, R1 and R2, and free centrioles, C1-4. Note the accessory structure on R1 in the bottom panel, confirming that the central centriole is a mother centriole. Section sequence is indicated in the bottom right of each panel. Scale bar = 1 μm. Also see figure S1B-E.

The centriole number in OSNs lies between that of two well-studied states. A common state is for cells to have exactly two centrioles, with the older of the two serving as a basal body for a primary cilium. This older centriole is often referred to as the mother centriole and the newer centriole as the daughter centriole. The daughter centriole forms orthogonally to the mother centriole in G1/S of the cell cycle and is engaged to the mother until mitosis (Kuriyama and Borisy, 1981; Nigg and Holland, 2018). Upon passage through mitosis, it becomes disengaged, and in the ensuing cell cycle, it acts as a mother centriole upon which another new daughter centriole can form. In contrast, multiciliated epithelial cells have as many as hundreds of centrioles, each serving as a basal body for a motile cilium. In this state, cells exit the cell cycle and initiate a transcriptional program that facilitates this centriole amplification(Hoh et al., 2012; Kyrousi et al., 2015; Ma et al., 2014; Tan et al., 2013). Centriole amplification in multiciliated epithelial cells occurs by two means: 1) centriole growth from deuterosomes, structures that are specific to multiciliated epithelial cells, and 2) by growth of multiple daughter centrioles from each mother centriole, forming rosettes (Sorokin, 1968). Centriole rosettes are thought to contribute only a small percentage of the total number of centrioles in multiciliated epithelial cells (Al Jord et al., 2014). Cycling cells can also be induced to form centriole rosettes by overexpression of Polo-like kinase 4 (Plk4), a kinase required for centriole duplication, or by overexpression of certain other centriole duplication proteins (Arquint et al., 2012; Habedanck et al., 2005; Leidel et al., 2005; Tang et al., 2011; Vulprecht et al., 2012).

Interestingly, centrioles in the olfactory epithelium were previously described to be arranged in a rosette-like array in some cells (Cuschieri and Bannister, 1975). However, the nature of these centriole rosettes and their possible relationship to OSN formation have not been investigated. Our work builds upon these findings by describing a role for rosettes in centriole amplification, identifying developmental timing of centriole amplification, and presenting a possible mechanism for driving centriole amplification in the mouse olfactory epithelium.

## Results

To better define the range of centriole number in OSNs in our preparations, we counted centrioles in OSNs from mice expressing eGFP-centrin2, a marker of the centriole (Bangs et al., 2015). In nasal septa from adult mice, OSNs had an average of 15.7 centrioles per cell, with wide variation around the mean (6 to 37 centrioles/cell, see **Figure 3F**) but no apparent trend across the anterior-posterior axis. This number of centrioles is similar to previous reports of cilium and centriole number in OSNs (Challis et al., 2015; Uytingco et al., 2019).

We next considered the potential means by which cells amplify centriole number during differentiation from stem cells to OSNs. Centrioles in the olfactory epithelium were previously described to be arranged in a rosette-like array in some cells (Cuschieri and Bannister, 1975). To assess the presence and role of centriole rosettes in the olfactory epithelium, we visualized centrioles by transmission electron microscopy (TEM) and fluorescence microscopy in both adult and embryonic tissue. First, ultrathin sections were made from dissected olfactory turbinates taken from adult mice and examined by TEM. We observed dendritic knob structures with multiple centrioles, typical of OSNs (**Figure 1A**), as well as horizontal basal cells with centriole pairs and primary cilia (not shown). Near the basal lamina, where OSN progenitor cells are typically found, we found cells with centriole rosettes (**Figure 1B**). Next, we determined whether centriole rosettes were present in cryosections of embryonic (E12.5) olfactory epithelia from mice expressing eGFP-centrin2 (**Figure 1C, S1A**). Rosettes were apparent as clusters of eGFP-centrin2 foci with the expected dimensions. Note that the cell shown in the inset has two rosettes, consistent with rosette formation on both preexisting centrioles. In addition, we found that rosette-bearing cells were positive for the neuronal marker β-tubulin III, confirming that these cells were committed to a neuronal cell fate (**Figure 1C**). Our results suggest that centrioles are amplified by rosettes in both adult and embryonic olfactory epithelium.

By observing olfactory epithelia of adult mice by TEM and embryonic mice by fluorescence microscopy, we also found cells that had centrioles in addition to two rosettes. In olfactory epithelia from adult mice, we found cells with two rosettes, as well as free centrioles by TEM (**Figure 1D, S1B-E**). Similarly, puncta of eGFP-centrin2 were observed near rosettes in embryonic olfactory epithelia by fluorescence microscopy (see **Figure 1C, S1A**). Whether free centrioles formed by detaching from a centriole rosette or free of a parental centriole (i.e. de novo) requires further investigation.

Next, we next asked whether centriole amplification can occur in cycling cells or only in non-dividing differentiated cells, using fluorescence microscopy in adult and embryonic olfactory epithelium. We used stage-specific markers to assess centriole amplification in cells in different stages of the cell cycle. In the olfactory epithelium of wild-type adult mice, some cells with nuclear PCNA, a marker for S phase, had centriole rosettes (**Figure S2A**). To determine whether cells which amplify centrioles in S phase proceed into mitosis, we probed for phospho-H3, a marker for mitosis, and confirmed the mitotic state by presence of condensed DNA. In the olfactory epithelium of wild-type mice, many mitotic cells had clusters of centrioles (**Figure S2B**), and in instances in which both spindle poles could be imaged, both poles had amplified centrioles (**Figure 2A**). Similarly, we identified newly-forming sister cells with condensed chromatin and amplified centrioles in embryonic olfactory epithelium at E12.5 (**Figure 2B**). It was not possible to precisely count centrioles in most mitotic cells due to limitations of imaging, particularly in adult olfactory epithelium sections. Instead, we applied a quantitative fluorescence method, measuring the total area of centrin fluorescence signal **(Figure 2C**). This method was calibrated on hTERT RPE-1 cells with and without Plk4 overexpression to generate centriole rosettes (**Figure S2C**). In the olfactory epithelium many of the mitotic cells had area measurements consistent with centriole clusters (rosettes +/− extra centrioles) rather than centriole pairs (**Figure 2C**). These results demonstrate that OSN precursors are able to divide after centriole amplification in both adult and embryonic olfactory epithelium, and that both sister cells of a division can receive an amplified collection of centrioles.

**Fig. 2.**
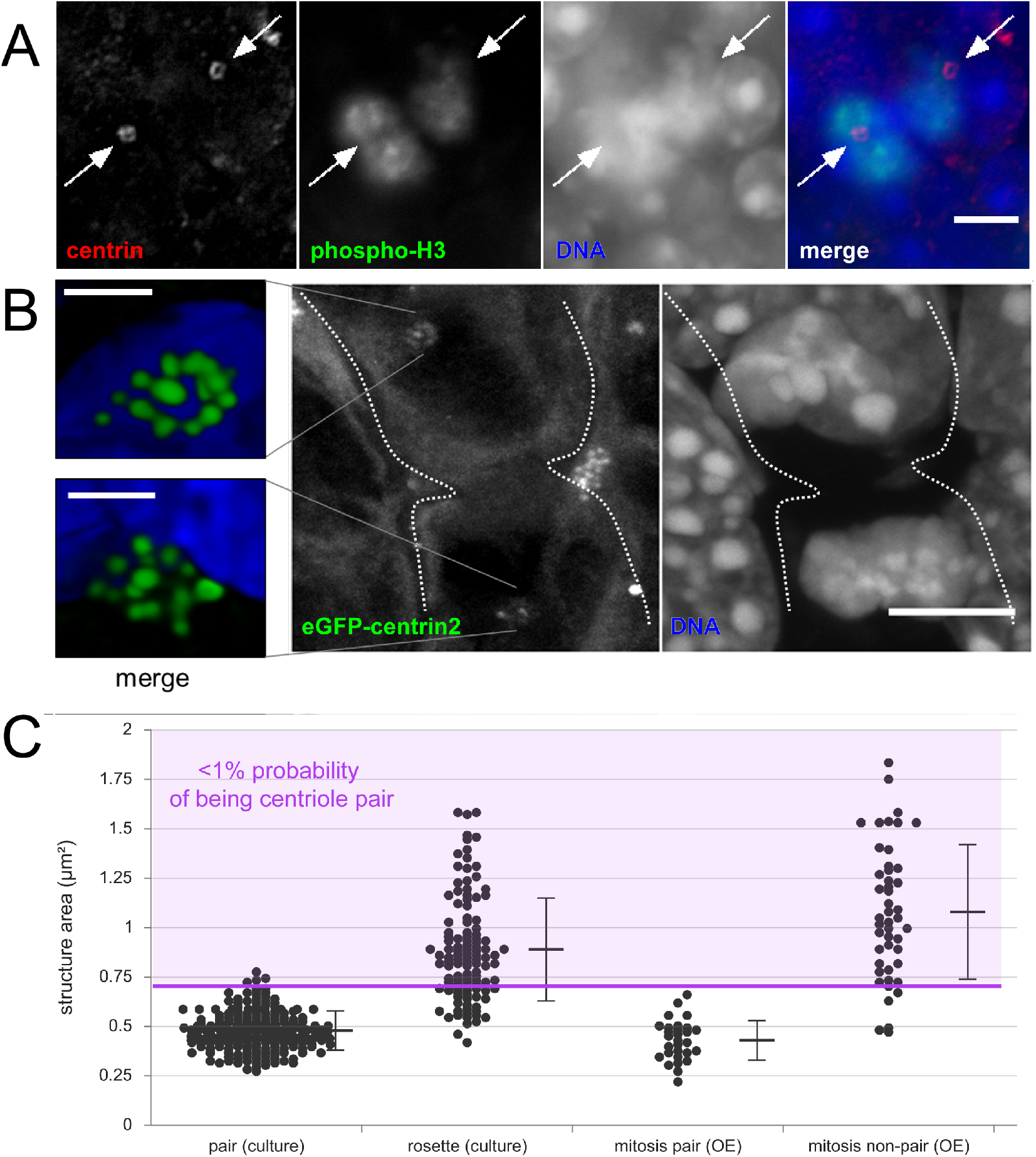
Division of cells with amplified centrioles. (A) Single optical section image of immunofluorescence in olfactory epithelium from a wild-type adult mouse. Phospho-H3 marks a cell in a late stage of mitosis. White arrows denote clusters of centrioles on opposite sides of the dividing cell. Scale bar = 5 μm. (B) Maximum projection image of fluorescence in olfactory epithelium from a mouse at embryonic stage E12.5. The mouse expresses eGFP-centrin2 to mark centrioles as well as Arl13b-mCherry (not shown). The insets show deconvolved images of centriole clusters in each newly-forming daughter cell. Dotted lines mark approximate cell boundaries. Scale bar = 5 μm. Inset scale bars = 1 μm. (C) Analysis of eGFP-centrin2 fluorescence area in mitotic cells in the olfactory epithelium. The pair (culture) column (N=3, n=208) shows measurements of centriole pairs in RPE-1 cells, which were used to set a threshold of 0.7085 μm^2^ (purple line), above which area measurements have □1% probability of belonging to the centriole pairs data set. The rosette (culture) column (N=3, n=115) shows measurements of centriole rosettes in cells overexpressing Plk4, 73.0% of which are above the threshold. The mitosis pair (OE) column (N=5, n=29) shows measurements of centriole pairs in adult olfactory epithelium, all of which fall below the threshold. The mitosis non-pair (OE) column (N=5, n=46) shows measurements of centriole structures which could not be definitively classified as pairs, 87.2% are above the threshold, Error bars = standard deviation. Also see figure S2.

To determine whether there is a gene expression program that might drive rosette amplification in the OSN lineage, we conducted a secondary analysis of an existing single-cell RNA sequencing (scRNAseq) data set. Fletcher et al. (2017) sequenced cells from dissociated mouse olfactory epithelium and grouped cells into distinct cell states by Slingshot analysis (Street et al., 2018). Analysis of a suite of genes associated with cell cycle progression showed that progenitors known as globose basal cells (GBCs) and early immediate neuronal precursors (INPs) are likely mitotically active (Fletcher et al., 2017). We specifically examined genes known to be upregulated in association with DNA synthesis, including *Rrm2*, which encodes ribonucleotide reductase 2 (Thelander and Berg, 1986). We found that the mRNA for many of these genes was abundant only in cells in the GBC and INP1 states, consistent with these being mitotically active (**Figure 3A**). Next, we analyzed the scRNAseq data for the expression pattern of genes encoding proteins associated with centriole duplication. Remarkably, the mRNA for Plk4 was strongly elevated in INP1s and INP2s, and less so in GBCs (**Figure 3B**). Plk4 effects centriole formation in conjunction with a binding partner, Stil (Arquint et al., 2012; Vulprecht et al., 2012). We found that the mRNA for Stil was also elevated in INP1s and INP2s (**Figure 3B, S3B**). The mRNA for Cep152, another binding partner of Plk4, also followed this pattern (**Figure S3A**), whereas those for most other centriole-associated genes did not. Given that upregulation of Plk4 or Stil mRNA drives centriole rosette formation in cell culture, we hypothesized that elevated Plk4 and Stil might drive centriole amplification in early INPs.

**Fig. 3.**
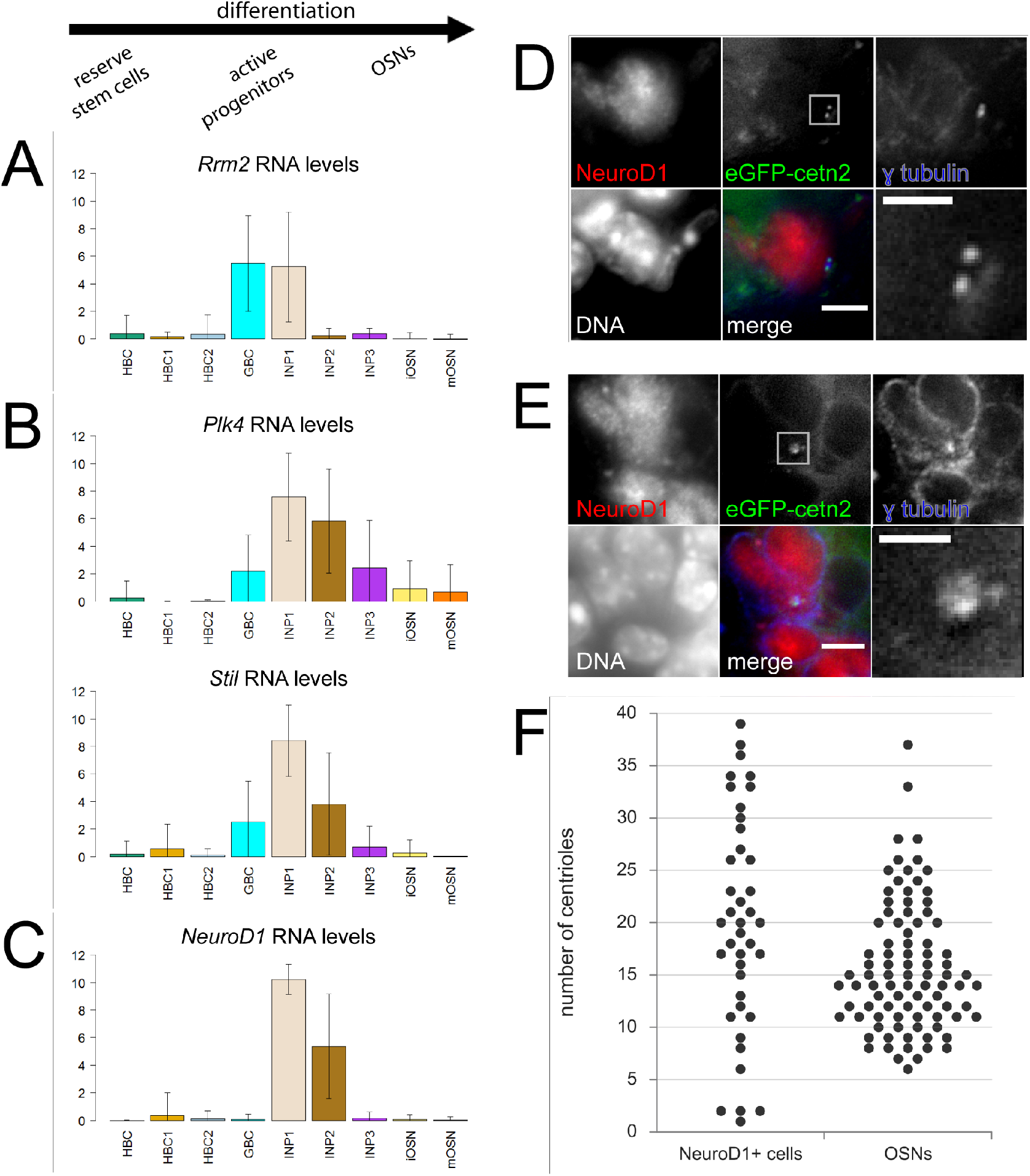
*Plk4* and *Stil* RNA levels and centriole number in early immediate neuronal precursors in the olfactory epithelium. (A-C) Secondary analysis of an existing single cell RNA sequencing data set from Fletcher et al. (2017) compares RNA levels for specific genes across cell types in the olfactory epithelium. The vertical axis shows average log_2_(normalized RNA counts). The horizontal axis shows cell groups in the lineage order as determined by Fletcher et al. (A) RNA levels for ribonucleotide reductase molecule 2 (*Rrm2*), a gene specific to DNA synthesis in S phase. (B) RNA levels for genes that drive centriole formation, polo-like kinase 4 (*Plk4*) and SCL/Tall interrupting locus gene (*Stil*). (C) RNA levels for *NeuroDI*, a transcription factor marking early immediate neuronal precursor cells. Error bars = standard deviation. (D-E) Fluorescence images of olfactory epithelium from adult mice expressing eGFP-centrin2 and Arll3b-mCherry. Cryo-sections were stained with antibodies against NeuroDI and y tubulin and with DAPI to mark DNA. (D) A NeuroDI-positive cell with two centrioles. (E) A NeuroDI-positive cell with non-pair centriole fluorescence. Scale bars = 5 μm. Inset scale bars = 2 μm. (F) Comparison of centriole counts in different cell types from olfactory epithelium of adult mice expressing eGFP-centrin2 and Arll3b-mCherry. Centrioles in NeuroDI-positive cells were counted in dissociated olfactory epithelia (N=2 mice, n=4O cells). Centrioles in OSNs were counted by en face imaging of the apical surface of septum olfactory epithelia (N=6 mice, n=9O cells). Note the small population of NeuroDI-positive cells with unamplified centrioles. Also see figure S3

To determine if elevated Plk4 and Stil mRNA levels correlate with the timing of centriole amplification, we used NeuroD1 as a marker of developmental timing within the differentiation pathway for OSNs. NeuroD1 is a transcription factor specifically upregulated in early INPs (**Figure 3C**) (Packard et al., 2011). We identified OSN progenitors in sections of adult olfactory epithelium by their localization near the basal lamina and presence of nuclear NeuroD1 immunofluorescence signal. We found examples of cells with two centrioles and cells with on-pair centrioles within the NeuroD1-positive progenitor population (**Figure 3D, 3E**). We used cells dissociated from olfactory epithelia of adult mice expressing eGFP-centrin2 to quantify centriole number in NeuroD1-positive precursors (**Figure 3F, S3D**). We compared these counts to the number of centrioles per OSN imaged in septa from adult mice expressing eGFP-centrin2. As in the sections, we found two groups amongst the NeuroD1-positive cells: a minority (n=4) of cells that had only one or two visible centrioles, suggesting that they had not yet amplified centriole number, and a majority (n=36) that had many more centrioles per cell (6 to 39 centrioles per cell). This distribution of centriole numbers suggests that centrioles are amplified in NeuroD1-positive cells, which also have high levels of Plk4 and Stil mRNA (**Figure S3C**). Together, our data support a model in which elevated Plk4 and Stil drive amplification via centriole rosettes in the olfactory epithelium.

## Discussion

Olfactory sensory neurons in mammals have a configuration of centrioles and cilia that distinguishes them from most other cells. We have found the centrioles in OSNs can be amplified from the progenitor cell’s centrosome via centriole rosettes prior to cell division, and that this is correlated with increased expression of the centriole duplication proteins Plk4 and Stil (**Figure 4**).

**Fig. 4.**
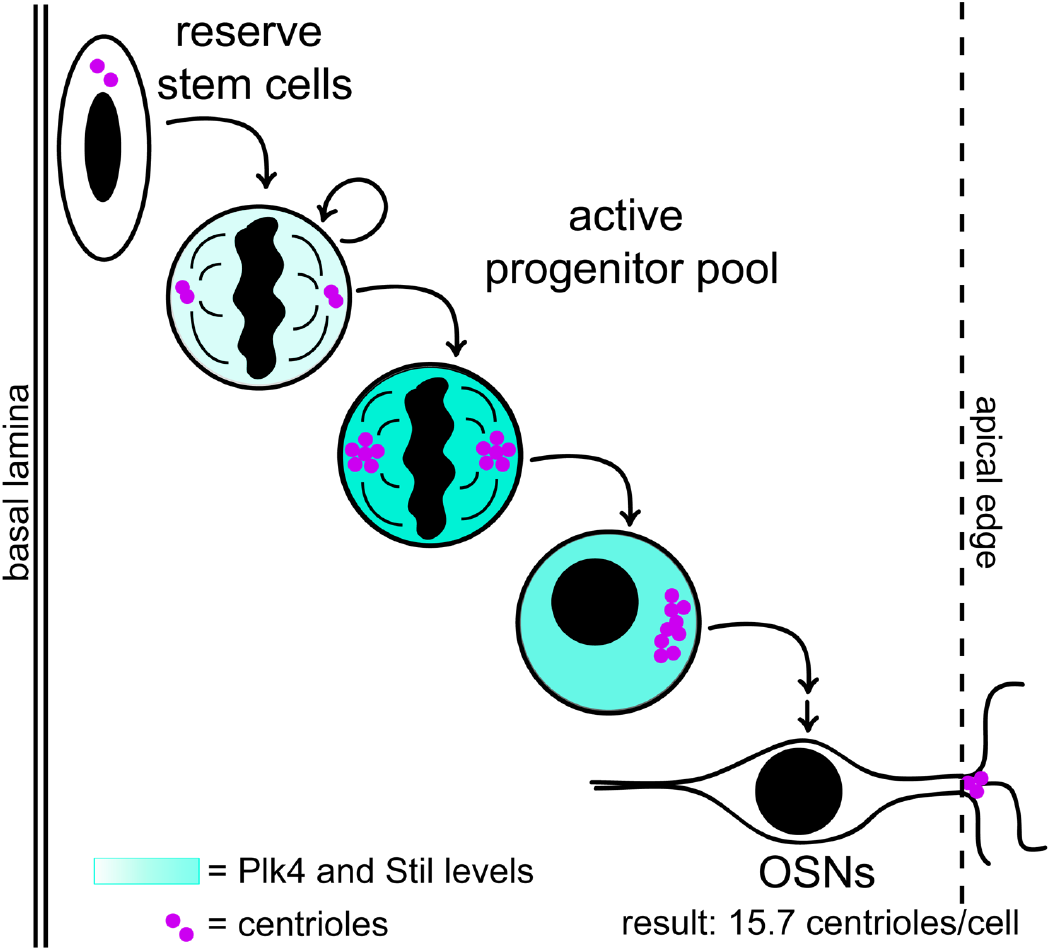
Visual summary of centriole amplification in the olfactory epithelium.

The number of centrioles in OSNs, an order of magnitude separated from most well-studied cell types, prompted us to ask how the observed number is achieved by cells. Our results show that centriole rosettes form on both of the pre-existing centrioles in progenitor cells in the olfactory epithelium and that both daughter cells receive an amplified set of centrioles, presumably one rosette each. If daughter centrioles in rosettes are engaged orthogonally to the mother as they are in cycling cells, then we estimate that the maximum total number of centrioles made by a rosette to be approximately nine (eight daughters plus one mother centriole), limited by the surface area around the base of the mother centriole. This is consistent with the number of daughter centrioles in rosettes observed in a single plane by TEM (see **Figure 1B**). However, the distribution of centriole numbers in OSNs shows that most have more than nine centrioles, thus, inheriting a single rosette of centrioles is insufficient to explain the observed number.

Within the constraints imposed by rosette size, there are several possible ways that OSNs might achieve the desired number of centrioles. One possibility is that more centrioles can be formed in a single amplification event. If centriole formation is not limited to the area around the base of the mother centriole, either by disengagement of daughter centrioles or by de novo synthesis, then many more centrioles can form in a single amplification event. Our observation of free centrioles in cells with rosettes supports this possibility, although our data do not distinguish between disengagement and de novo synthesis. We note that the presence of free centrioles in addition to rosettes would allow the possibility of asymmetric segregation of centrioles during mitosis, which would yield cells with centriole numbers near the extremes of the observed distribution. The second possibility for how OSNs achieve the desired number of centrioles is that additional centriole amplification might occur in subsequent cell cycles or after the final cell division in OSN differentiation. There is precedence for the latter in multiciliated epithelial cells, the only other widely-studied example of centriole amplification in vertebrates, where cells only amplify centrioles post-division.

Our data show that centriole amplification can occur in mitotically active progenitors of the olfactory epithelium. This is surprising because division with amplified centrioles is usually considered to be detrimental due to chromosome missegregation (Ganem et al., 2009; Silkworth et al., 2009). However, several features of the process might mitigate the potential problem of mitosis with amplified centrioles. First, many of the amplified centrioles are contained within rosette structures that likely function as single organizing centers during spindle formation, based on the single mother centriole within each rosette (**Figure S1B-E**). Indeed, we found that each rosette in this case had only a single focus of γ-tubulin (**Figure S2A**). This is similar to what appears to occur in Viviparus spermatogenesis, during which cells with rosettes go through meiosis (Gall, 1961; Pollister and Pollister, 1943). Cosenza et al. (2017) showed that the fidelity of mitosis in cells with overexpression-induced rosettes is sensitive to asymmetry in the number of daughter centrioles per rosette. It is unknown whether this phenomenon plays a role in the OSN lineage, and this would require live imaging of mitoses in the olfactory epithelium to resolve. Second, newly-formed free centrioles would not have undergone the centriole-to-centrosome conversion that would promote their ability to form a spindle pole in the mitosis immediately following their formation (Wang et al., 2011). Even if the free centrioles were capable of microtubule nucleation, known mechanisms could enforce bipolar spindle formation, for example by HSET-dependent centriole clustering (Kwon et al., 2008).

We showed that the transcripts for key proteins in centriole duplication, Plk4 and Stil, are transiently upregulated during OSN differentiation. This raises the question of whether this upregulation is sufficient to coordinate the formation of centriole rosettes. In multiciliated epithelial cells, *Plk4* and other centriole-associated genes are upregulated to drive centriole amplification via deuterosomes and rosettes during differentiation (Hoh et al., 2012). OSN differentiation closely resembles that of multiciliated epithelial cells, except that no deuterosomes form in OSN differentiation. This is likely because *Deup1*, a gene specific to and necessary for deuterosome formation (KlosDehring et al., 2013), is not expressed in the OSN lineage (data not shown); hence, only rosettes form. Besides *Plk4* and *Stil*, the only other centriole-associated gene that followed the same pattern of increased RNA levels in early INP cells was *Cep152*. Interestingly, Cep152 protein is known to be necessary for anchoring Plk4 and Stil at the mother centriole (Hatch et al., 2010), but increased Cep152 has not been associated with rosette formation. Additionally, how Plk4 and Stil transcription or RNA half-life is increased in early INPs remains unclear.

An important potential application of our work is to the development of neuron regeneration therapies. The olfactory epithelium contains the most accessible pool of adult neuronal stem cells, making it an attractive source for repairing diseased and damaged neurons in adults. Outside of the olfactory epithelium, amplification of centrioles would not be desirable, and our work highlights a window within OSN differentiation during which cells might be used to generate other types of neurons. Fitting with our findings of when centriole amplification likely occurs, GBCs are thought to give rise to cells both with amplified centrioles (OSNs) and without amplified centrioles (microvillar cells), whereas INPs ultimately give rise to only OSNs (Fletcher et al., 2017). Thus, if OSN progenitors are used to generate other types of neurons, upstream progenitors may produce more robust differentiation into neuronal cell types, as centriole number has repercussions for neuronal cytoskeletal structure and for formation of primary cilia, which are important for signaling in many neuronal cell types (Green and Mykytyn, 2014; Guemez-Gamboa et al., 2014).

In summary, our work highlights a system in which centriole amplification and cell division occur as a normal part of development and organ maintenance. These findings outline an important window for therapeutic potential of olfactory stem cells and also reveal more information about the basic biology of centriole formation.

## Materials and Methods

### Mice and cells used

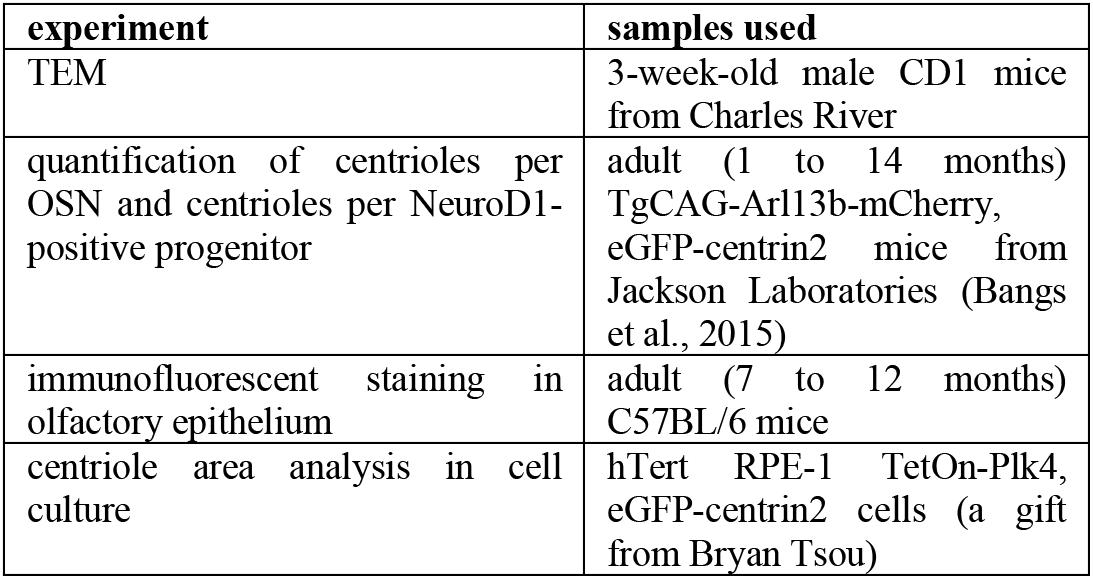

### Antibodies used

Primary antibodies used for immunofluorescent staining are listed in the table below. AlexaFluor-conjugated secondary antibodies (Thermo-Fisher) were diluted 1:1000.

| target | source | dilution | treatment |
| --- | --- | --- | --- |
| $\beta$ tubulin III | BioLegend, clone TuJ1 | 1:2000 | none needed |
| $\gamma$ tubulin | Sigma-Aldrich, clone GTU88 | 1:1000 | none needed |
| centrin (all isotypes) | EMD Millipore, clone 20H5 | 1:5000 | 10 minute pretreatment in 0.5% (w/v) sodium dodecyl sulfate in PBS |
| phospho-H3 | Cell Signaling, #53348T | 1:200 | none needed |
| GFP | Invitrogen, #A-11120 | 1:1000 | none needed |
| Sass6 | Santa Cruz Biotech, #91.390.21 | 1 μg/mL | requires methanol fixation of cells at -20°C |
| NeuroD1 | Proteintech, #12081-1-AP | 1:100 | 10 minute pretreatment in 0.5% (w/v) sodium dodecyl sulfate in PBS, overnight incubation in primary antibody |

### Transmission electron microscopy

Mice were euthanized by CO_2_ in accordance with Stanford’s APLAC guidelines. Facial bones were removed in a dish of cold Tyrode’s solution (140mM NaCl, 5mM KCl, 10mM HEPES, 1mM CaCl_2_, 1mM MgCl_2_, 1mM sodium pyruvate, 10mM glucose in ddH2O), as in other reports (Dunston et al., 2013), and turbinate scrolls were mechanically separated from septa. Epithelia from turbinate scrolls were removed mechanically and fixed immediately in a solution of 2% glutaraldehyde and 4% PFA in 0.1M Na cacodylate buffer for 3 to 4 hours at 4°C. Samples were then rotated in a 1% solution of OsO_4_ for 1 hour at room temperature, washed four times gently in water, then rotated in a 1% solution of uranyl acetate overnight at 4°C. Samples were then dehydrated in a graded ethanol series (30%, 50%, 70%, 95%, 100%, 100%) for 15 to 20 minutes per step, rotating at room temperature. Samples were washed twice for 10 minutes each in propylene oxide (PO), then embedded through a graded PO:EMBED resin series (2:1 for 1 hour, 1:1 for 1 hour, 1:2 overnight). Samples were then rotated in pure EPON with lids open for 5 hours to evaporate remaining PO before embedding in molds at 50°C for 4 days. Semi-thin sections were taken and imaged on a dissecting scope to find samples in the correct orientation. 80 nm sections were treated with uranyl acetate and mounted on grids before imaging on a JEOL JEM-1400 transmission electron microscope. Samples were prepared from two separate animals, and rosettes were observed in both.

### Immunofluorescent staining of cryosections

Olfactory epithelia were dissected as described above. Whole olfactory epithelia, turbinate epithelia, or E12.5 embryo heads were fixed immediately in 4% PFA in PBS at 4°C for 3 to 24 hours. Samples were then washed in phosphate buffered saline (PBS) and stored at 4°C. Before mounting, samples were equilibrated in 1 to 5mL of 30% sucrose solution in water for a minimum of 12 hours at 4°C. Samples were embedded in OCT compound (Sakura Tissue-Tek) on dry ice and stored at −80°C. Embedded samples were sectioned at 8 to 14 μm on a Leica cryostat and adhered to charged slides by drying at room temperature for approximately 1 hour. Slides were stored with drying pearls (Thermo Fisher) at −80°C and thawed under desiccation no more than twice. Samples were pretreated as needed (see antibody summary chart) and rehydrated and blocked for 0.5 to 4 hours in 5% milk in 0.1% Triton-x 100 that had been spun in a tabletop centrifuge to pellet un-dissolved milk particles. Slides were incubated in primary antibody for approximately 3 hours, washed in PBS, incubated in secondary antibody for approximately 1 hour, washed in PBS, incubated in DAPI for 1 to 5 minutes, washed in PBS, and mounted in MOWIOL. Embryonic samples were imaged on a spinning disk confocal microscope, and adult samples were imaged on a Zeiss inverted widefield microscope using MicroManager (Edelstein et al., 2010). Images were processed in FIJI (Schindelin et al., 2012). Images from the widefield microscope were deconvolved using the Iterative Deconvolve plug-in (Dougherty, 2012) and theoretically-generated point spread functions (Diffraction PSF 3D). For images with high background, contrast in the representative images was adjusted uniformly across the image such that the area outside of cells was black and areas of high signal were just below saturation. Each immunofluorescent staining procedure was performed at least three times (N≥3) with samples taken from at least two separate animals.

### Analysis of centriole structure area

To quantify the area of centriole structures, samples were imaged on a Zeiss inverted widefield microscope. For proof of concept, centrioles from RPE-1 cells with immunofluorescent staining (see procedure below) were imaged in z-stacks with 0.5 μm steps to include all centrioles in the field of view (N=3 replicates from cell seeding through staining). 30 images of each condition were taken and processed in FIJI. Z-stacks were converted into a maximum projection image, and the green channel was deconvolved using the Iterative Deconvolve plug-in (Dougherty, 2012). Engaged structures were selected based on the presence of anti-Sass6 immunofluorescence signal between adjacent anti-GFP puncta, and rosettes were defined as structures with at least three anti-GFP puncta. We measured the area of anti-GFP fluorescence in structures meeting these criteria and normalized all measurements such that the average area of centriole pairs was exactly 2. The normalized fluorescence area had approximately a 1:1 ratio with actual centriole number (0.9208), demonstrating that it is an appropriate approximation for centriole number (**Figure S2C**). Actual area measurements were approximately the dimensions expected for two centrioles, modeled as the projection of two overlapping spheres (analysis not shown). Next, we used this method to assess centriole structures from the olfactory epithelium (**Figure 3C**). We estimated the probability density distribution of centriole pair areas to be a Gaussian function. We used this distribution to estimate a cutoff area (0.7085 μm^2^) above which a structure has less than 1% probability of belonging to the centriole pairs’ dataset. As a proof of concept, 73.0% of rosettes measured in cell culture were above this cutoff. For olfactory epithelia, single-plane images were used for analysis shown here, though similar results were obtained with z-stacks (data not shown). We applied the cutoff to mitotic cells of the olfactory epithelium because, in contrast to S-phase cells, mitotic cells’ centrioles separate in preparation for spindle formation, reducing overlap and making structures more amenable to measurement. Images of anti-centrin immunofluorescence signal in mitotic cells in cryosections from adult mice were deconvolved using the Iterative Deconvolve plug-in, and area was measured by outlining puncta in the anti-centrin channel. Images in which centriole pairs were clearly visible were categorized as such. All other images were categorized as “non-pair” structures. Immunofluorescent staining was performed five times, and all mitotic cells centrioles that could be imaged were included. 87.2% of area measurements in this group fell above the cutoff. Probability calculations and modeling of area projections were performed in R. Dot plots were generated with Statistika (Weissgerber et al., 2017).

### Overexpression of Plk4 in cell culture

RPE-1 cells were cultured in DMEM/F-12 (Corning #MT-10-092-CV) with 10% cosmic calf serum (GE Healthcare #SH30087.04) and periodically tested by PCR for mycoplasma contamination. Stock cultures were selected by hygromycin B (Thermo Fisher #10687010) prior to the start of experiments. Cells were seeded to be 70-80% confluent at the start of the experiment. S-phase arrest was initiated by incubating cells in 2mM thymidine. After 24 hours, media was replaced with new media containing 1μg/mL doxycycline and 2mM thymidine to induce overexpression, or with 2mM thymidine and no doxycycline for the control condition. After 24 additional hours, cells were washed in PBS, fixed for 20 minutes in methanol at −20°C, washed again in PBS, and stored at 4°C.

### Immunofluorescent staining of RPE-1 cells

Cells cultured on poly-L-lysine-coated coverslips were fixed in methanol at −20°C for 20 minutes and washed in PBS. Samples were blocked for a minimum of 30 minutes in 5% dry milk in 0.1% Triton-x 100 that had been spun to pellet undissolved milk particles. Samples were then washed 3 times in PBS, incubated with primary antibodies for 1 to 2 hours at room temperature, washed 3 times in PBS, incubated with secondary antibodies for 0.5 to 2 hours, washed 3 times in PBS, incubated with DAPI for 5 minutes, washed 3 times in PBS, and mounted in MOWIOL. Cells were imaged on a Zeiss inverted widefield microscope with MicroManager (Edelstein et al., 2010). Images were processed in FIJI.

### Secondary analysis of scRNAseq data

Single-cell RNAseq data were obtained from Fletcher et al. (2017). Cells were pooled for average RNA levels based on categories determined by Fletcher et al. Analyses were performed in R (code available upon request).

### Quantification of centrioles in OSNs

Samples were dissected as described above, except that dissections were performed in PBS. Septa were immediately transferred to 4% paraformaldehyde (PFA) in PBS and fixed for 3 to 24 hours at 4°C. Septa were washed and stored in PBS at 4°C. For imaging, septa were mounted in a chamber of double-sided tape on glass slides with SlowFade Gold mountant (Invitrogen) and high precision 1.5 weight coverslips (Deckglässer) sealed with nail polish. Samples were imaged on a Leica SP8 scanning confocal microscope. For each sample, five fields of view were spaced approximately evenly along the anterior-posterior axis of the olfactory epithelium. Within each field of view, the cell at the center of each quadrant of the field was imaged such that the z-stack included all centrioles within the dendritic knob. The lowest-quality image from each field of view was excluded from the analysis, giving 15 cells per animal (N=6 animals, n=90 cells total). Individual dendrites were identified by their tightly-clustered centrioles (data not shown). Image stacks were processed by semi-automated detection in the program Imaris x64 9.2.1 using the Surfaces function and separating touching objects by seed points of 0.3 μm diameter. Dot plots were generated using Statistika (Weissgerber et al., 2017).

### Olfactory epithelium dissociation

The turbinate region of olfactory epithelia was dissected in cold Tyrode’s solution, as described above. Samples were incubated in 1 to 2mL of 0.25% trypsin (Thermo-Fisher, #MT-25-053-CI) and minced with a feather scalpel periodically, between incubations at 37°C, for 10 to 15 minutes in total. Trypsin was inactivated by adding 10 mL of DMEM with 10% serum. Samples were poured over a 40 μm cell strainer to remove bone fragments and other large debris. Samples were spun at 800g for 5 minutes to pellet, washed in PBS, and spun again. Samples were resuspended and fixed in 4% PFA in PBS overnight at 4°C, then washed and stored in PBS at 4°C.

### Quantification of centriole number in progenitor cells

Dissociated samples (N=2) were stained to identify NeuroD1-positive progenitors by first spinning at 8000rpm for 2 minutes in a tabletop centrifuge to remove PBS, then pretreating by resuspending in 0.5% (w/v) SDS in water for 1 minute. Cells were spun to remove SDS, then resuspended and blocked for 0.5 to 1 hour at 4°C in a solution of 5% dry milk in 0.1% Triton-x 100 that had been spun to remove un-dissolved milk particles. Samples were washed in PBS, incubated in primary antibody overnight at 4°C, washed in PBS, incubated in secondary antibody for 30 minutes at room temperature, washed in PBS, incubated in DAPI solution for 2 minutes, washed in PBS, then resuspended in MOWIOL. Samples were mounted with 1.5 weight coverslips, and centrioles were imaged (n=20 cells per sample, n=40 total) and counted by the same method as OSN centrioles, described above.

## End Matter

### Author Contributions and Notes

Conceptualization, writing, and funding acquisition: K.C. (ORCiD: 0000-0002-0517-2421) and T.S. (ORCiD: 0000-0002-0671-6582); Investigation: K.C.; Supervision and project management: T.S.

The authors declare no competing interests.

## Acknowledgments

We thank John Ngai, Russell Fletcher, and Diya Das for helpful suggestions and feedback on the secondary analysis of their scRNAseq data, Eszter Vladar for helpful discussions, Emily Kolenbrander Ho for feedback on the manuscript, Mary Mirvis for assistance with image analysis, and John Perrino and Stanford’s Cell Sciences Imaging Facility for help with TEM. EM imaging was supported, in part, by ARRA Award Number 1S10RR026780-01 from the National Center for Research Resources (NCRR). This project was supported by the CMB Training Grant from NIGMS of the National Institutes of Health under award number T32GM007276, by the National Science Foundation Graduate Research Fellowship under Grant No. D–G16E5 6518, and by the NIGMS of the National Institutes of Health under award number 1R35GM130286-01. We thank Stanford Bio-X for access to microscopy resources, and members of the Stearns lab for helpful feedback and suggestions.

## Supplemental Information

**Supplemental figure S1.**
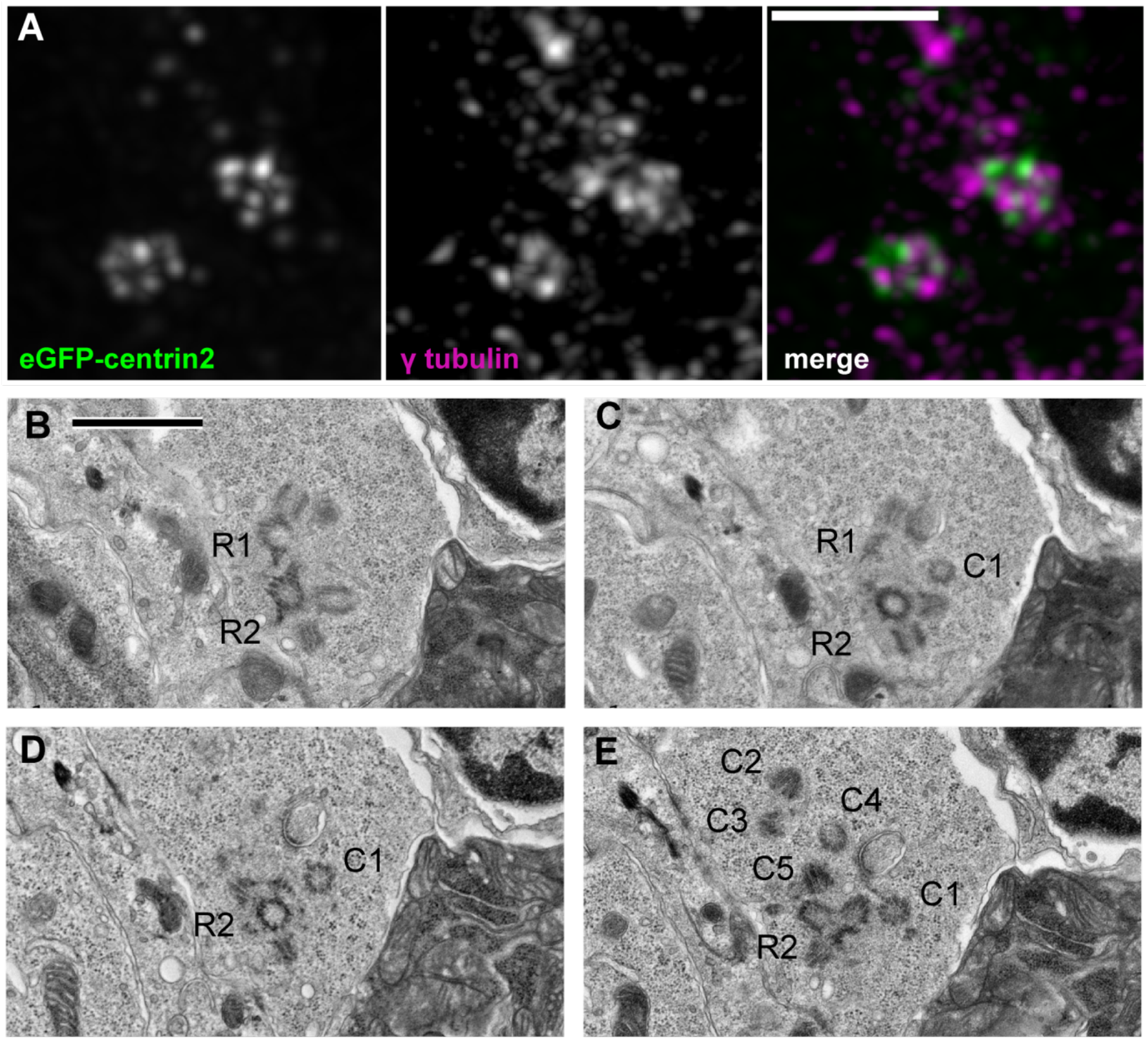
Co-existence of engaged and non-engaged centrioles. (A) Inset from a maximum projection fluorescence image of embryonic olfactory epithelium at E12.5 in mice expressing eGFP-centrin2 to mark centrioles. Deconvolved images show two rosette-like centriole clusters and separate puncta positive for centriole markers eGFP-centrin2 and γ tubulin. Scale bar = 2 μm. (B-E) TEM images from serial sections of olfactory epithelium from a wild-type adult mouse. R1, R2 denote centriole rosettes, identified by morphology and accessory structures on the mother centriole. C1-5 denote centrioles not associated with rosettes. Note that both mother centrioles in panel A have accessory structures. Scale bar = 1 μm.

**Supplemental figure S2.**
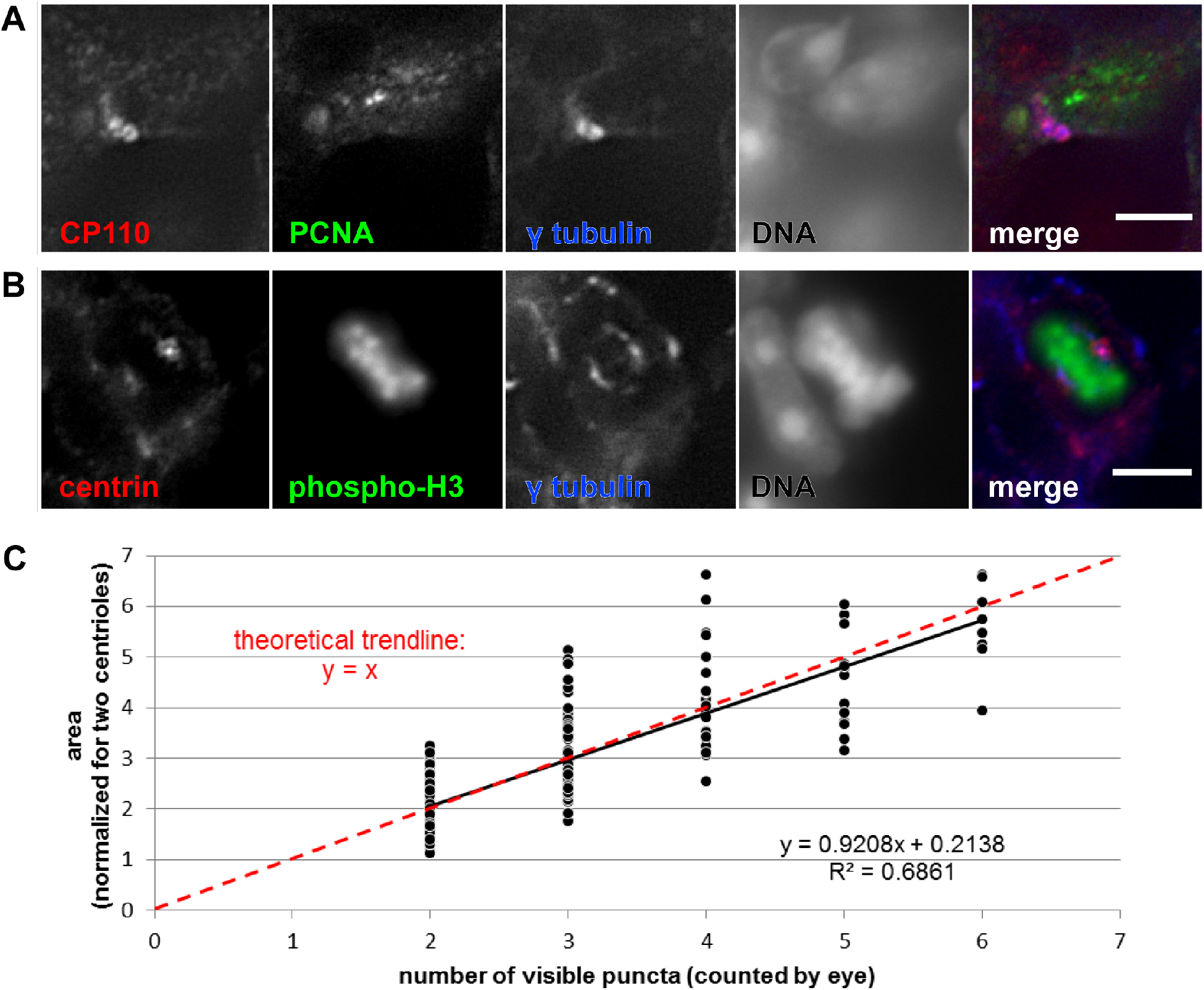
Division of cells with amplified centrioles in the olfactory epithelium. (A) Immunofluorescence in cryosections of olfactory epithelium from a wild-type adult mouse. Punctate nuclear PCNA marks a cell in S phase, whereas nearby nuclei are PCNA-negative. CP110 marks the distal ends of centrioles, γ tubulin marks centrosomes, and DAPI marks DNA. In this single optical section, daughter centrioles are visible as rings around γ tubulin foci, consistent with rosette formation. Scale bar = 5 μm. (B) Immunofluorescence in cryosections of olfactory epithelium from a wild-type adult mouse. Phospho-H3 marks a cell in mitosis, and DAPI shows DNA condensed and aligned in metaphase. Centrin marks the distal ends of centrioles. In this single optical section, centriole clusters are shown at the spindle poles, marked by γ tubulin. Scale bar = 5 μm. (C) Plot of anti-GFP fluorescence area against centriole number in cell culture. Immunofluorescence images were taken of hTert RPE-1 TetON-Plk4, eGFP-centrin2 cells with and without doxycycline induction. Anti-GFP fluorescence area of Sass6-positive structures was measured, and puncta were counted by eye. A line of best fit was generated in Microsoft Excel. The slope of the line is 0.9208, showing an approximately linear relationship between centrin fluorescence area and centriole number.

**Supplemental figure S3.**
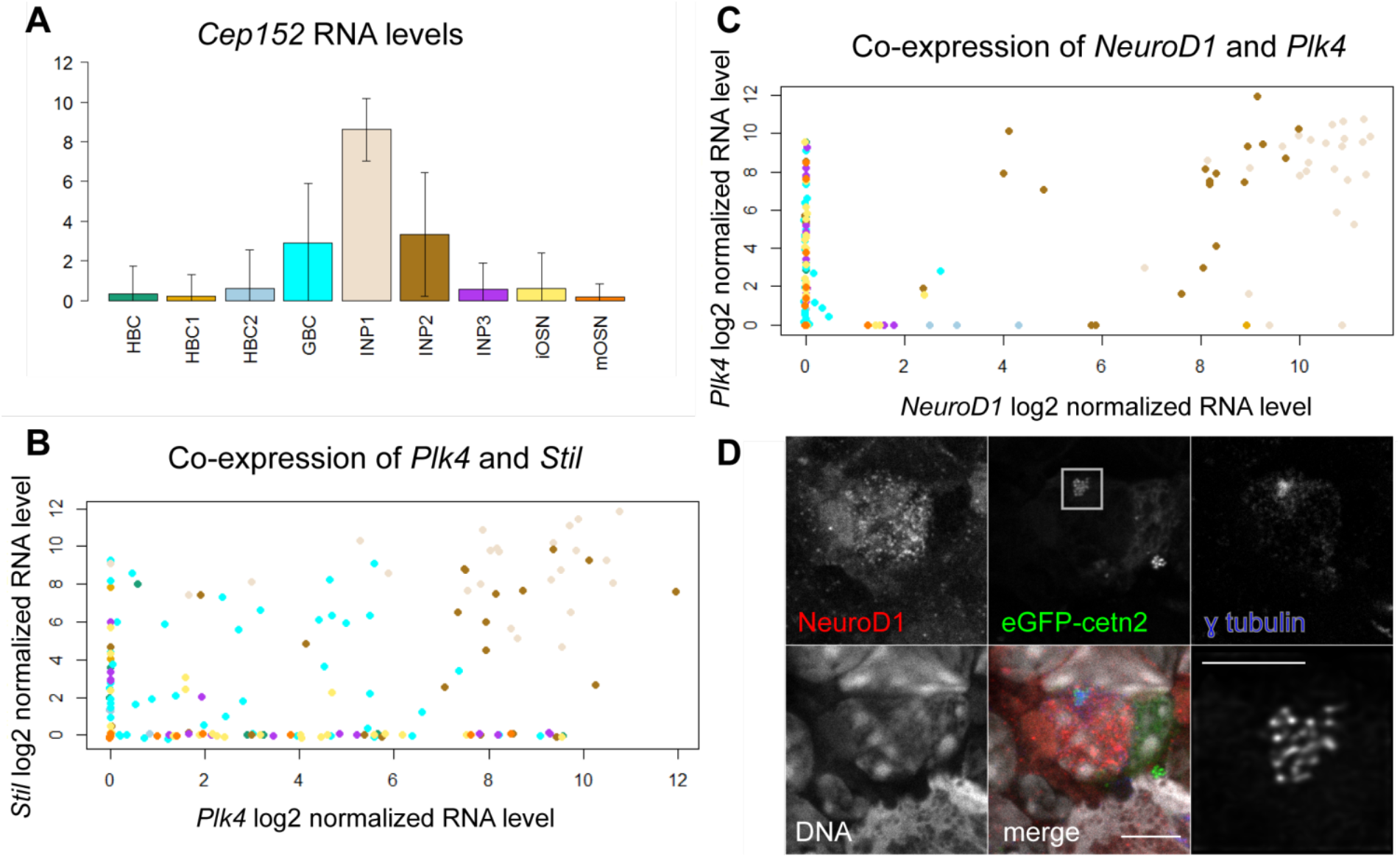
RNA levels in scRNAseq data and images of a NeuroD1-positive cell. (A) Secondary analysis of an existing single cell RNA sequencing data set from Fletcher et al. (2017) compares RNA levels for specific genes across cell types in the olfactory epithelium. The vertical axis shows log2(normalized RNA counts). Plot shows RNA levels for *Cep152*, a gene that interacts with *Plk4* in centriole formation. (B-C) Secondary analysis of data from Fletcher et al., 2017 shows co-expression of genes. Each dot represents a single cell. Both axes show log2(normalized RNA counts) (B) Plot shows RNA levels for *Plk4* and *Stil*, centriole-associated genes known to drive rosette formation in cell culture. Points in the upper right corner are cells that express both genes at high levels. These are INP1 and INP2 cells (see A for color coding). (C) Plot shows RNA levels for *NeuroD1*, a transcription factor that marks INP1 and INP2 cells, and *Plk4*. Points in the upper right corner are cells that express both genes. These show that individual INP1/2 cells express high levels of *Plk4*. (D) A fluorescence image of a NeuroD1-positive cell in dissociated olfactory epithelium. Note that this image includes other nuclei that are NeuroD1-negative. The inset shows a deconvolved image of centrioles, marked by eGFP-centrin2. Scale bar = 5 μm. Inset scale bar = 2 μm.

